# The evolutionary history of topological variations in the CPA/AT superfamily

**DOI:** 10.1101/2020.12.13.422607

**Authors:** Sudha Govindarajan, Claudio Bassot, John Lamb, Nanjiang Shu, Yan Huang, Arne Elofsson

## Abstract

CPA/AT transporters consist of two structurally and evolutionarily related inverted repeat units, each of them with one core and one scaffold subdomain. During evolution, these families have undergone substantial changes in structure, topology and function. Central to the function of the transporters is the existence of two non-canonical helices that are involved in the transport process. In different families, two different types of these helices have been identified, reentrant and broken. Here, we use an integrated topology annotation method to identify novel topologies in the families. It combines topology prediction, similarity to families with known structure, and the difference in positively charged residues present in inside and outside loops in alternative topological models. We identified families with diverse topologies containing broken or reentrant helix. We classified all families based on 3 distinct evolutionary groups that each share a structurally similar C-terminal repeat unit newly termed as “Fold-types”. Using the evolutionary relationship between families we propose topological transitions including, a transition between broken and reentrant helices, complete change of orientation, changes in the number of scaffold helices and even in some rare cases, losses of core helices. The evolutionary history of the repeat units shows gene duplication and repeat shuffling events to result in these extensive topology variations. The novel structure-based classification, together with supporting structural models and other information, is presented in a searchable database, CPAfold (cpafold.bioinfo.se). Our comprehensive study of topology variations within the CPA superfamily provides better insight about their structure and evolution.

## Introduction

Proteins from the CPA/AT (Monovalent cation/proton antiporter /Anion transporter) superfamily transport a variety of ions, amino acids and other charged compounds [1–4] (Table S1). Due to their functional importance, CPA/AT transporters are ubiquitously present in all the three kingdoms of life [5–8]. In humans, CPA/AT transporters are associated with pathological conditions such as intestinal bile acid malabsorption, ischemic and reperfusion injury, heart failure and cancer [9,10]. Therefore, these transporters serve as important drug targets. [11,12].

All known CPA/AT structures consist of two inverted symmetric ***repeat units*** (Figure 1a). These repeats are essential to enable the different conformational states that are necessary for the transport mechanism [13–16]. Each of the repeat units can be further divided into two structurally distinct parts, the ***scaffold*** and the ***core subdomains*** (Figure 1a) *[7]*. The two N and C scaffold and core subdomains come together in structure to give rise to one scaffold and one core domain (Figure 1b).

**Figure 1:**
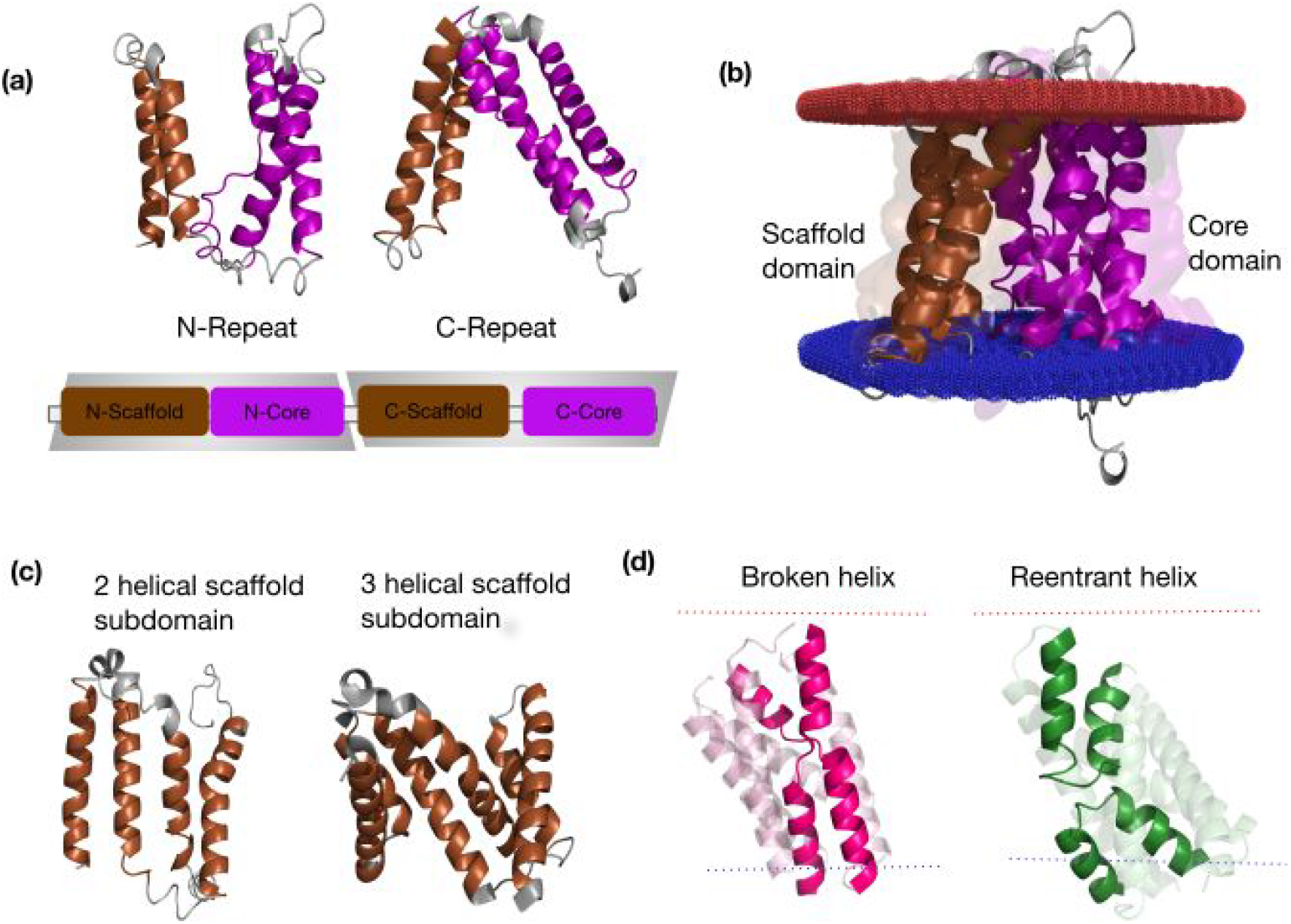
General structure of CPA/AT transporters: (a) The figure shows the sodium bile acid symporter (PDB id:4n7w). N-terminal repeat is followed by C-terminal repeat (In inverted orientation) in sequence space. Each repeat unit is composed of a scaffold and a core subdomain. The repeats are shaded in grey. (b) In structure space, both the N-scaffold and C-scaffold subdomain come together to form a scaffold domain. Similarly, the N-core and C-core subdomain come together to form the core domain. The lipid bilayer is coloured red and blue to denote outside and inside respectively. (c) Scaffold subdomain can be 2 or 3 helices long. (d) Core domain can be broken (coloured in dark pink) or reentrant (Coloured in green).

The scaffold domain is involved in dimerisation and also interacts with the core domain (Figure 1c). The interface between the two domains forms an aqueous cavity where the substrate binds. The core domains generally consist of 6 transmembrane helices, with the middle helix of the core subdomains being a non-canonical, broken or reentrant, helix. The broken helix is a transmembrane helix that crosses the membrane with a discontinuity in the alpha-helix, forming a loop within the membrane. In contrast, the reentrant helix does not cross the membrane and enter and exit from the same side (Figure 1d). Therefore, the topology of the following helices has to differ to accommodate the different location of the C-terminal end of the reentrant/broken helices. The non-canonical helices contain a polar non-helical part in the centre of the membrane region capable of binding/transporting ions [17] and transferred to the other side of the membrane with elevator mechanism [18].

Proteins with known structure are available only for 5 families in the superfamily. It varies starting from 10 transmembrane helices (TM) in the SBF family [19] to 12 TM in Na_H_Antiport_1 and OAD_beta [20,21], and 13 TM of Na_H_Exchanger and 2HCT [7], [22].

The CPA/AT clan is classified differently in different protein databases. In Pfam, the CPA/AT transporters belong to the “CPA/AT clan” [5]. However, it should be noted that the sodium-citrate symporter (2HCT, PF03390) is not included in the CPA/AT clan, although it is evolutionarily, structurally and functionally related to the other members. In the TCDB database [23], the members are split into the CPA- and BART-superfamilies [24], while in OPM [25] there exists only one superfamily, the “Monovalent cation-proton antiporter superfamily”. Further, ECOD [26] and CATH [27] classifies members into two different superfamilies. The discrepancy suggests a need to investigate the relationship between CPA/AT transporter families from a structural and evolutionary perspective.

A systematic study of topological diversity within the CPA/AT superfamily is still missing. Therefore, we analysed the evolutionary mechanism of the repeat units and identified three distinct conserved repeat units that we name “Fold types”. All data can be found in a searchable database, CPAfold (cpafold.bioinfo.se).

## Results

### Expanding the structural coverage of the CPA/AT superfamily by topology annotation

We updated and filtered the Pfam clan (CPA/AT) and named it “CPA/AT superfamily”, containing at the end 15 Pfam families. In addition to the original families in the Pfam clan, two families, Abrb (PF05145) and 2HCT (PF03390) were added to the superfamily. Further, families with reliable alignments with other families are critical for reliable topology annotation. Therefore, four families, OAD_beta, (PF03977) lys_export (PF03956), sbt_1 (PF05982) and *LrgB* (PF04172) were excluded as they are only distantly related (See methods section).

As described in the methods section, the integrated pipeline (Figure 2) annotates the families with topologies using a combination of topology prediction, similarity to families with known structure, and the difference in positively charged residues present in inside and outside loops in different topological models. Among the 15 families/subfamilies, we identify proteins with novel topologies containing 9 to 14 transmembrane helices. Eight families have a broken helix in their core domain, while seven families have a reentrant helix (Table 1). The annotations agree with earlier experimental topology annotations [28] [29] [30].

**Figure 2:**
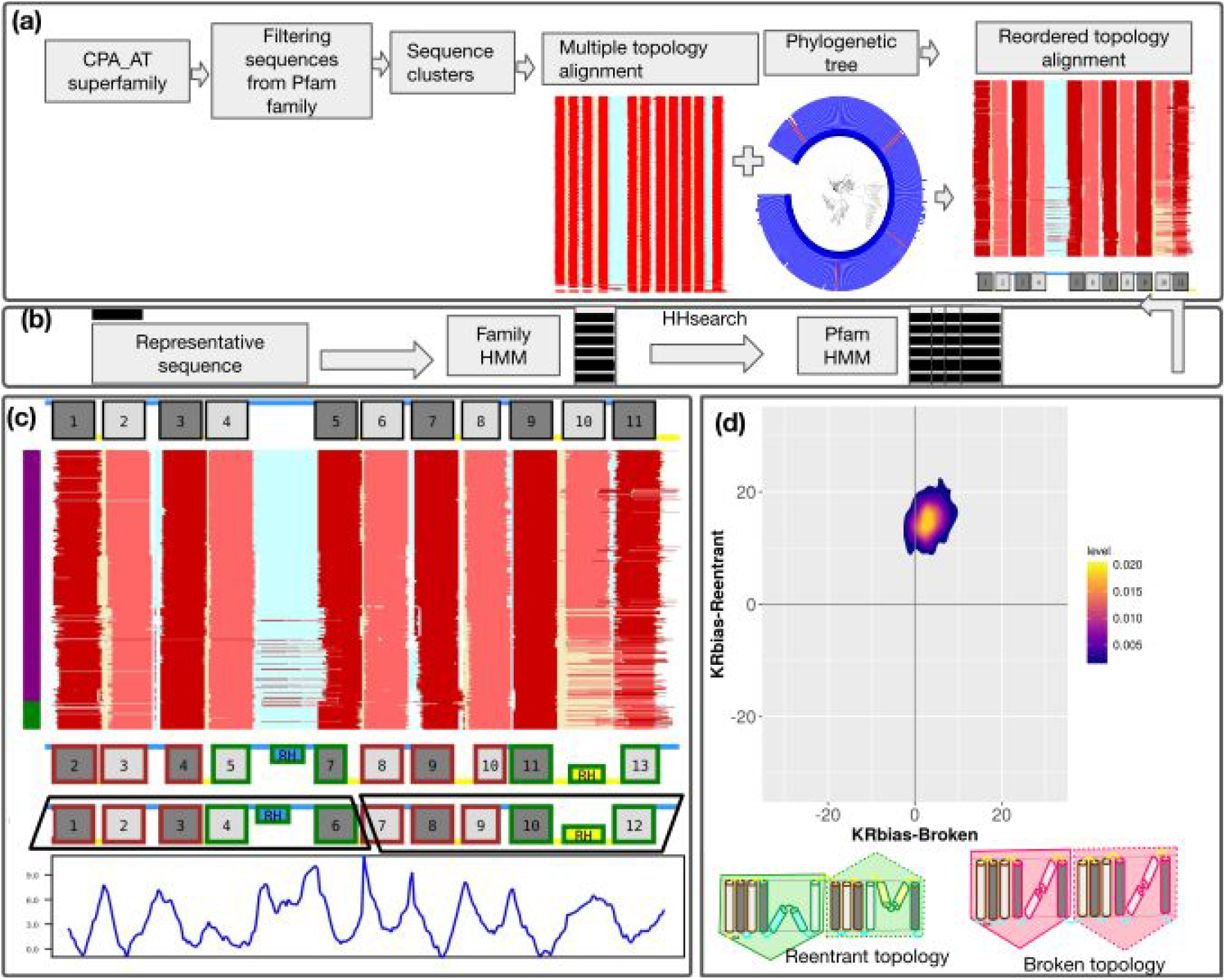
An integrated pipeline to annotate the topology of CPA/AT transporters: Steps involved in the pipeline are shown. (a) Identification of Pfam subfamilies, with a unique topology and assignment of initial topologies. (b) Improved classification of CPA/AT superfamily (c) Generation of final topology and annotation of core and scaffold subdomains are shown in the DUF819 family. The figure shows a reordered topology alignment. The TM helices (in-out) and TM helices (out-in) are coloured dark red/grey and light red/grey, respectively. Reentrant helices (in-in) and (out-out) are coloured yellow and blue respectively. The inside and outside loops are coloured yellow and blue respectively. The vertical bar is coloured based on the taxonomy of the sequences (Bacteria: Purple, Archaea:Dark blue Eukaryotes: Green). Scaffold subdomains and reentrant core subdomains are coloured brown and green, respectively. N and C repeats are shown as black trapezoids. ΔG values are obtained for the representative sequence and are plotted to the aligned residues in the representative sequence. (d) Validation of broken/reentrant transporters by using the positive inside rule for the DUF819 family. The mean KR bias value is higher for the reentrant model than the broken model.The KR-bias plot is shown as a 2D scatter plot. The correct reentrant topology model shows inverted repeats.

**Table 1:**
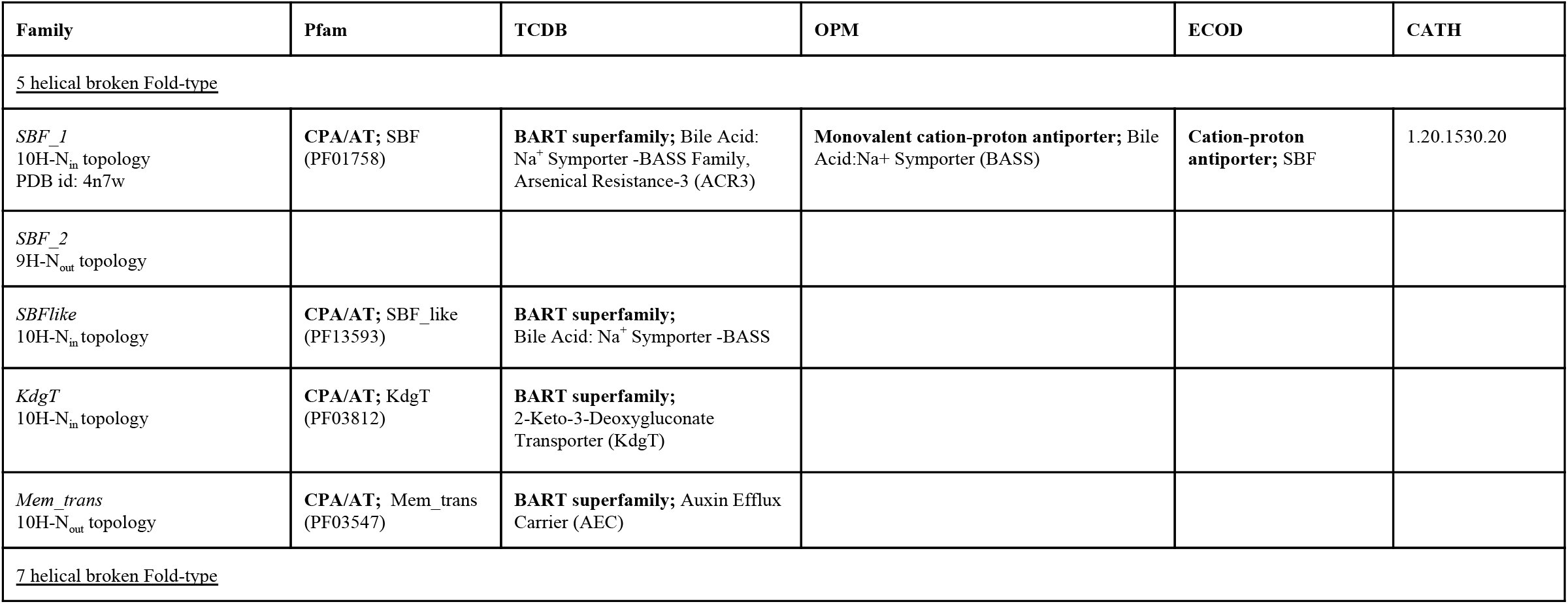
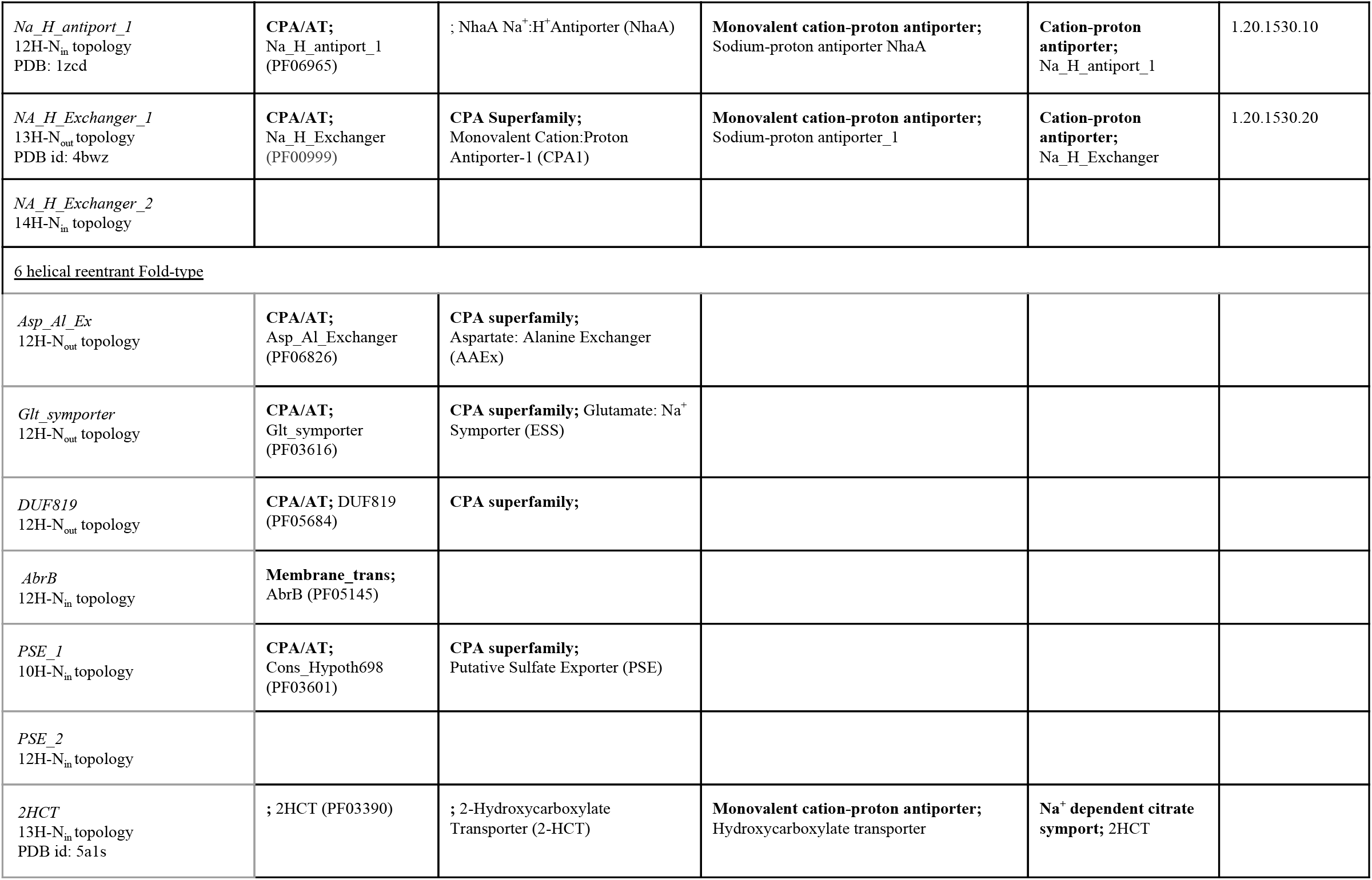
Annotation of topology and subdomains for families/subfamilies in the CPA/AT superfamily: The orientation (N_in_, N_out_) describes the location of the N-termini of the protein. Comparison of classification of families within the CPA/AT transporter superfamily (as defined here) and in different databases are shown. Superfamily/clan classifications are shown in bold, followed by family classifications.

### CPA/AT transporters cluster into three “Fold-types”

A dendrogram made from sensitive MSA-MSA pairwise alignments of the separate N and C repeat units depicts the evolutionary relationship of the families (Figure 3). In the dendrogram, three distinct clusters can be identified - each with the same number of helices in the C-terminal repeat, as the topology of the N-terminal repeat unit varies in some families. We define these as three separate “Fold-types”; (1) The 5-helical-broken fold-type, (2) the 7-helical-broken fold-type, and (3) the 6-helical-reentrant fold-type.

**Figure 3:**
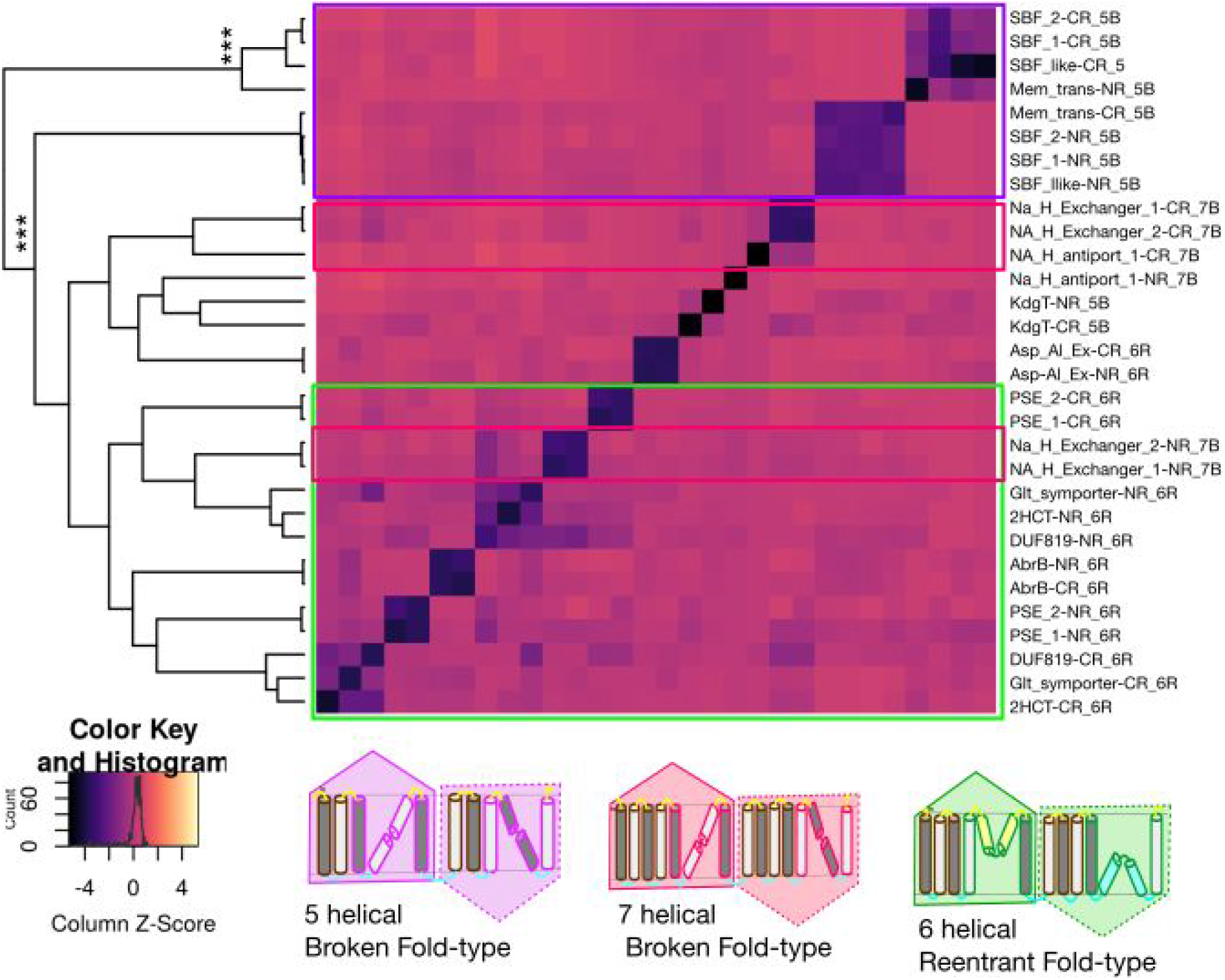
Repeat units are clustered based on conserved C-terminal repeat: Fold-types. Hierarchical clustering, and heatmaps obtained using E-values between N- and C-terminal repeats of different families. E-values (log) are converted to Z-scores (scaled column-wise) to generate the dendrogram. The clusters with *p*-value *greater than or equal to p=0.95 (***)* is represented as asterisks.

The clustering shows that the 7-helical-broken and 6-helical-reentrant fold-types are closest, while the 5-helical-broken is an outlier. The full-length transporters also cluster into three fold-types (Figure S1), and this clustering is statistically significant, see Figure S2 and S3). A maximum-likelihood tree also confirms the evolutionary relationship between the three fold types (Figure S4 and S5). However, the maximum likelihood trees suffer from low bootstrap values in a few nodes, due to high sequence divergence between families.

### Different types of topology variations and their evolution

The CPA/AT families are homologous but have different topologies. Therefore, during the evolutionary history of CPA/AT transporters, the topology has changed. The clustering of the repeat units of families suggests that several different types of evolutionary events have occurred (Figure S6). Below we will list the four necessary events.

#### 1) Broken-reentrant transporter transitions

The most notable topology transition between fold-types is the transition between the broken and reentrant helices (Figure S6). The transition does affect not only the helices themself but also all the following helices as they have to change their orientation. It also requires the gain/loss of a helix in the scaffold subdomains to maintain the inverted nature of repeat units. In Figure 4a and 4b, it can be seen that the gain/loss of a helix always occurs in the C-termini scaffold subdomain.

**Figure 4:**
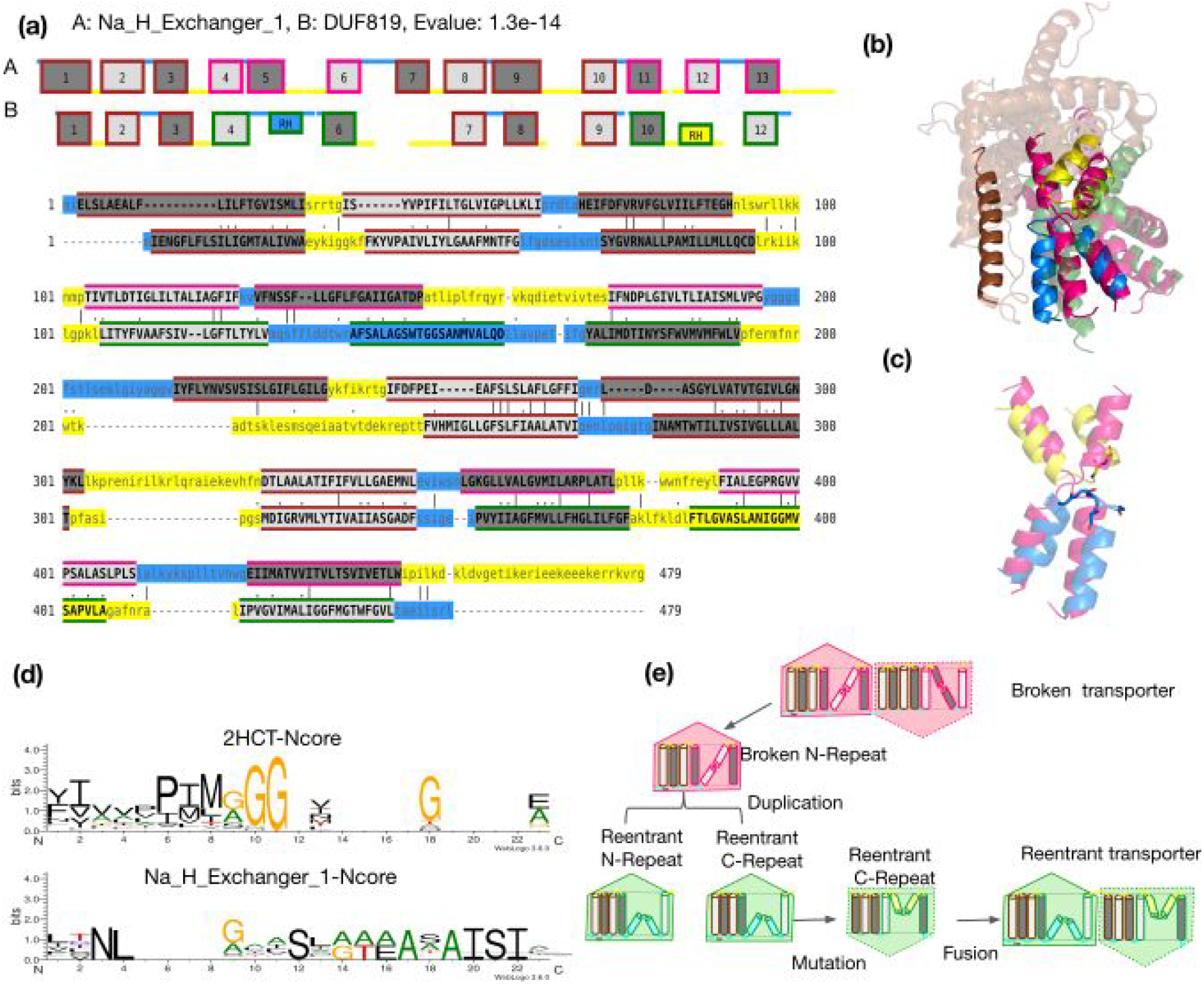
Broken-Reentrant transporter transitions. (a) Sequence and topology alignment between 7 helical broken transporter and 6 helical reentrant transporter.(b) Structure superposition of broken and reentrant transporter. The extra scaffold helix and the broken-reentrant transition are highlighted in bright colours. (c) A zoomed-in figure of broken (pink) and reentrant helix (yellow and blue) is shown.The glycines are shown in stick representation. (d) The aligned positions of the broken and reentrant N-core helix are represented as sequence motifs (e) Cartoon representation showing the events of duplication and mutation in reentrant transporters leading to the transition of broken-reentrant transporters.

The alignment shows that the central non-helical regions of the reentrant helices are enriched in glycine residues, whereas the broken helices are not (Figure 4c and 4d). Core sequence motifs for all the families are available in CPAfold database.The polar enrichment in the reentrant helices makes it less hydrophobic and more prone to bending than the broken helices. (Figure 4c and S7a).

It is well known that the cytoplasmic side of membrane proteins are enriched in positive residues (K and R) [31]. The transition from a broken to a reentrant region changes the orientation of the last helix in the core subdomain (Figure S7b). Therefore, the number of positively charged residues in the surrounding loops changes.

Anyhow, structural alignment also shows the difference in the packing of the helices in the broken and reentrant core subdomains, caused by the change in orientation of the third helix. However, overall the broken helical and reentrant core domains align well (Figure S7c).

The repeat units of different fold-types that appear to be closest are the N-terminal repeat of Na_H_Exchanger family (7-helical broken) and both N and C repeat units from DUF819 (6-helical reentrant), see Figure S8a and S8b. A model for the evolution of this structural transition could include (not necessarily in this order); (i) Mutations in N terminal repeat of the seven helical broken repeats leading to reentrant helix (ii) Duplication of the repeat unit, (iii) fusion of the repeats, and (iv) mutations to change the orientational preference of one of the repeats to form a functional reentrant transporter (Figure 4e). This shows a potential path for the closely related N-terminal repeat of the Na_H_Exchanger family (7-helical broken) to change into the DUF819 (6-helical reentrant).

#### 2) Changes in orientation in reentrant transporters

Families from the same fold-type can have opposite orientations (Figure 5a and S9). The repeat level clustering and alignments clearly show that AbrB, PSE_1, and PSE_2 families undergo orientation changes compared to all the other reentrant transporters in this superfamily (Table 1).

**Figure 5:**
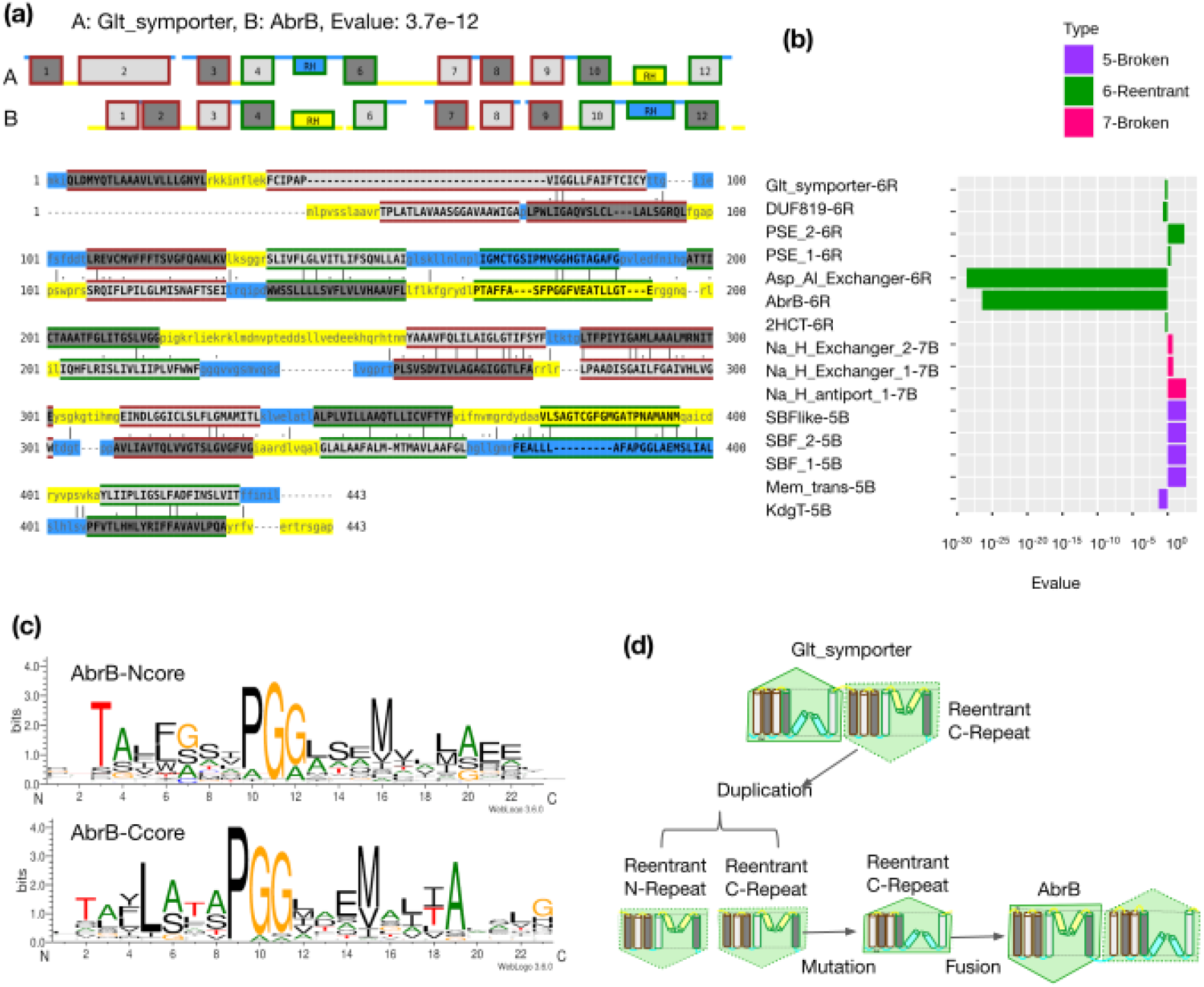
Change in orientation in reentrant transporters. (a) Sequence and topology alignment between reentrant transporters showing the change in orientation. (b) Sequence similarity between N and C repeats represented as E-values in different families of the superfamily (c) Reentrant N and C core helix motif (d) Cartoon representation showing the events of duplication from C-terminal repeat of reentrant transporter and subsequent internal duplication leading to change in orientation.

It is necessary to remember that all CPA/AT transporters have an internal symmetry, i.e. their origin is an internal duplication of these repeat units at some point of their evolutionary history. In most families, the internal duplication is more ancient than the divergence between protein families, as the N-terminal repeats from different families are more similar than repeats within the same protein family (i.e. between N-terminal repeat and C-terminal repeat). An exception to the general observation is the high similarity between the N- and C-terminal repeat units of the reentrant transporters, Asp-Al_Exchanger and AbrB families (Figure 5b). Due to the recent duplication, both the N- and C-terminal reentrant helices retain the signal of a glycine-proline motif (Figure 5c). This reentrant motif enriched with both glycines and prolines have been identified for the first time in CPA/AT superfamily. One scenario describing this evolutionary change starts from the C-terminal repeat of an ancestral Glt-symporter. This repeat unit is then internally duplicated and fused followed by mutations to change the orientation of the C-terminal-repeat, see (Figure 5d and S10). Asp_Al_exchanger also has a recent internal duplication in the N_in_,-region, i.e. it retains the same orientation as the other reentrant families. This can be explained by a duplication starting from an N-terminal repeat unit.

#### 3) Changes in orientation in broken transporters

Inversion of the orientation also occurs in the broken transporters. The Mem_trans family have an opposite orientation compared to all the other families of the same fold-type (eg. SBF_like) (Table 1). The N-terminal repeat of the SBF_like family is similar to the C-terminal-repeat of Mem_trans and vice versa (Figure 6a, 6b and S11). This reciprocal similarity between the repeat units shows that there has occurred a recent shuffling of the repeat units causing the change in orientation, see Figure 6c.

**Figure 6:**
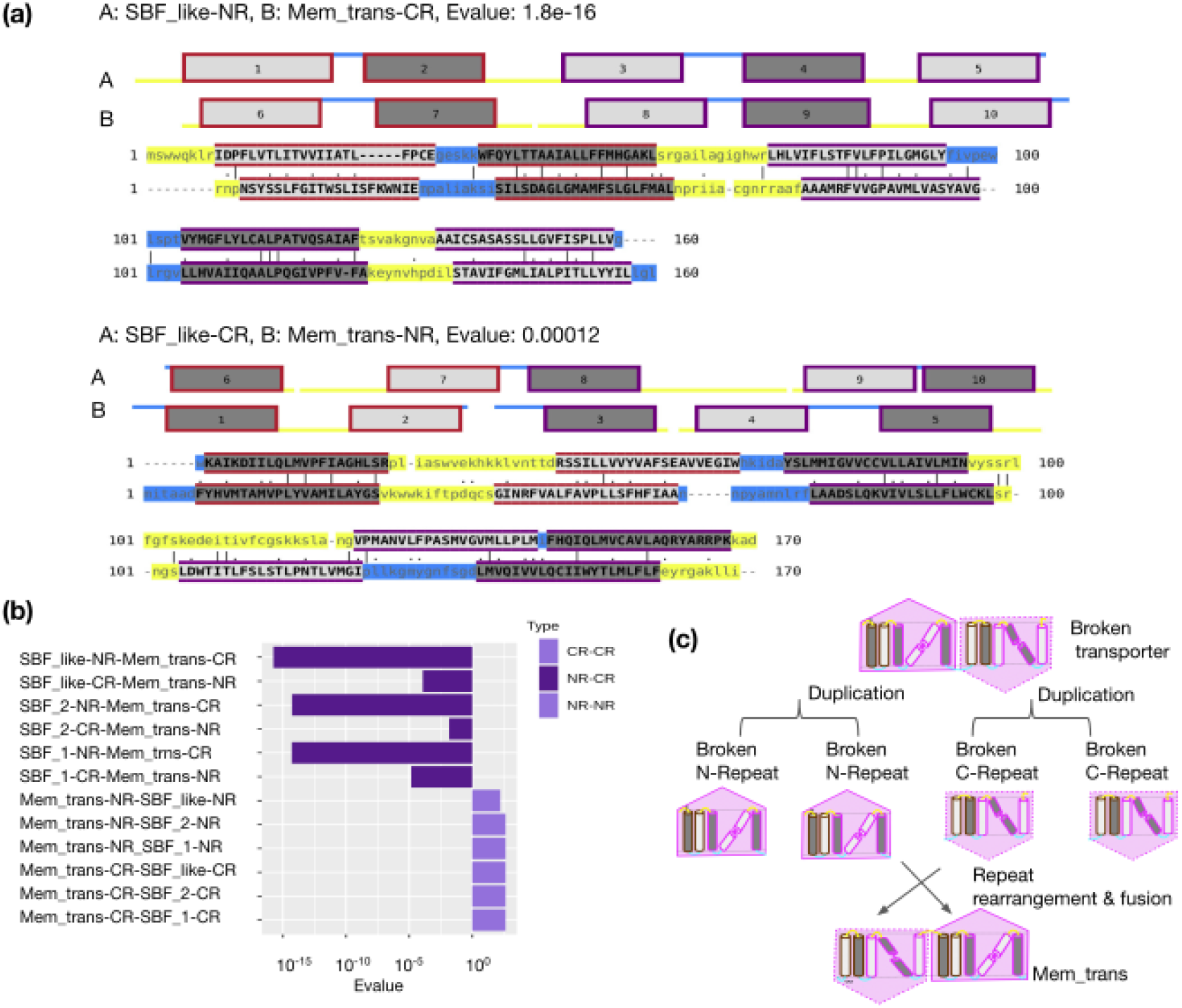
Change in orientation in broken transporters. (a) Sequence and topology alignments, which shows the high similarity between N-terminal and C-terminal repeats of SBF_like and Mem_trans families and vice versa. (b) Sequence similarity, represented as E-values, between N-N, C-C and N-C repeats of two different families in the 5-helical-broken fold-type are shown. (c) Cartoon representation showing events of shuffling of repeats leading to change in orientation.

#### 4) Gain/loss of Scaffold helix and rare loss of core helix

It is generally assumed that the topology is conserved within a family [32]. However, members of SBF, Na_H_Exchanger and PSE families have two distinct topologies. Therefore, these families were split into two subfamilies based on their unique topology (Table 1). Topology variations between subfamilies/families always show gain/loss of helices in the N-terminal scaffold subdomain (Figure S12 and S13). When switching fold types, variations of topology always occur in the C-terminal scaffold subdomain but may or may not occur in the N-terminal scaffold (Figure S14). We also observe that two helices have been lost in the C-terminal core subdomain of PSE_1(Figure S15), PSE_2 subfamilies.

## Discussion

The presence of non-canonical helices such as broken/reentrant helices makes it challenging to base topology prediction purely on automatic tools [33,34]. Instead, we found that the best way to identify broken and reentrant helix prediction was by aligning the families to known structures. We then use the “KR-bias” plots to confirm the predicted topologies (Figure 2d). Significantly, cases with a change in orientation are not evident from the alignment to known structures and need to be corrected. Also, when the unknown topology is longer than the known topology, topology predictions become essential.

We believe that the statistically significant clusters from hierarchical clustering are phylogenetically correct as hierarchical clustering is capable of capturing the evolutionary history of highly divergent superfamilies [35]. In general, our structure-based classification is consistent with the classification from TCDB [36,37]. In TCDB the 5-helical broken fold type is classified into the BART superfamily [24], while the CPA superfamily contains both the 6-helical reentrant and the 7-helical broken fold types [2].

Here, we propose a new structure-based classification scheme for the CPA/AT superfamily based on a conserved C-terminal repeat unit called “Fold-types”. This classification is different from the classical Protein fold-superfamily-family hierarchy in SCOP [38]. It shows a larger structural diversity than in most other protein superfamilies. The CPA/AT superfamily can be classified based on the decreasing order of structural and evolutionary relatedness into (1) Superfamily, (2) Fold-types, (3) Family and (4) Subfamily (Figure S16). Despite showing structural similarity in the C-terminal repeat, families belonging to the same fold-type might contain variations in the N-terminal repeat unit (Figure S12 and S13). We believe that this classification scheme could, in the future, be applied to other membrane proteins containing repeat units.

Some families, including Na_H_exchanger, SBF, PSE families have more than one topology. Presence of eukaryotes in these families pinpoints that topology variation could be more prevalent in eukaryotes than in bacteria or archaea, (See the CPAfold database, for species distribution for topologies in different families). Previous work suggests that alternative splicing can result in variations of topology. Functional isoforms exist in the Na_H_Exchanger and SBF families [39–41].

Variations of topology in the scaffold domain contribute to both structural and functional advantages for the transporters. In the case of transporters with a three helical scaffold subdomain, the first helix helps to dimerise the scaffold domain (Figure 1c). The dimerisation of the scaffold domain is important for the elevator mode of transport [7,42–44]. It is already known that asymmetry in domains results in functional specialisation of elevator type transporters [18]. The functional reason for the change in orientation in the Mem_trans family is to enable the transport of the plant hormone auxin out of the cell [45]. Enhanced stability and function are observed in proteins that have evolved by recent duplication [46]. Our analysis of CPA/AT transporters shows that repeat units are capable of rearrangements or shuffling in complicated ways during evolution, using similar mechanisms as in the evolution of multidomain proteins by domain rearrangements [47]. Our comprehensive computational study has helped to understand better the structure, function and evolution of the transporter superfamily.

### Experimental procedures

#### 1) Topology annotations of families and subfamilies

Our strategy to annotate topology and reclassify the Pfam CPA/AT clan into families/subfamilies involves the following five steps, also described in Figure 2.

I. Identification of Pfam subfamilies, each with a unique topology and assignment of initial topologies.
II. Improved classification of CPA/AT superfamily
III. Generating a final topology.
IV. Identification of core, scaffold subdomains and repeat units from the known structure.
V. Validation of Broken/reentrant type transporters by the positive inside rule.

##### i) Identification of Pfam subfamilies, each with a unique topology and assignment of initial topologies

We extracted reference proteome sequences from the 13 Pfam families in the CPA/AT clan [31,48]. Fragments, sequences with <75% Pfam domain coverage, and highly similar sequences (>90% identity) were excluded. The remaining sequences were clustered at 30% identity using blastclust [49] and aligned using Clustal Omega [50]. Topologies for all the members of the families were predicted using TOPCONS2 [51]. For families with a systematic variation in the predicted topologies, the MSA is then split based on the topology, and the families are split into subfamilies that each have a unique topology. This MSA was named as “Topology seed MSA”. It was noted that the topology predictions sometimes consistently missed predicting either one or both broken/reentrant helices. Therefore, the topologies annotated using the reordered topology alignments were labelled as “Initial topologies”. To assign correct topology to all families within the CPA/AT superfamily, we, therefore, had to use additional steps (Figure 2a).

##### ii) Improved classification of the CPA/AT superfamily

A representative sequence for each family was selected, In case of families with a known structure, this sequence was selected. In cases without a known structure, the top hit from a single search with HMMsearch [52] against Uniprot was used. The representative sequence of each family was searched against Uniclust30 [48] using the HHblits program [53] with default settings generating a representative or family MSA.

We used HHsearch version 3.2.0 [54] to find possible evolutionary relationships between the family MSA of the CPA/AT clan and other families in Pfam-A_v32.0 [5]. We wanted to search for new Pfam families that are not yet assigned to be part of CPA /AT superfamily. We defined the “CPA/AT superfamily” to only include Pfam families that have an E-value better than 10^−4^ with at least one family of the clan (except itself). New families that are added to the CPA/AT superfamily went through the previous step to annotate their initial topology and check if they have topology variations within the family (Figure 2b).

##### iii) Generating final topology

The next step was to identify the missing, broken/reentrant helices if any. The representative sequence of the family/subfamily was searched against the PDBmmCIF70_22_May database using HHsearch [54] to compare the “initial topology” of the family with the topology derived from the crystal structure. Missing transmembrane helices (reentrant or broken) in the representative sequence were inferred from the alignment to the known structure.

Predicted topology and the topology from the crystal structure were mapped onto the pairwise MSA-MSA alignment to obtain a topology alignment. Transmembrane helices were considered to be aligned when at least five residues of both helices are aligned, as used before [55]. Otherwise, it is a, TM helix aligned to gap regions, TM helix aligned to inside/outside loops, TM helix aligned to signal peptide. The type is chosen based on the dominating composition in the segment of the TM helix that is aligned. Missing helices in the representative sequence were inferred when the TM helix in the topology with known structure was aligned to loops in the representative sequence. Since we have structural templates for both broken and reentrant type transporter, classification is based on the type of transporter with a known structure with best hit (lowest E-value) (Table S2). Alignment with known structures also helped to correct incorrect topology prediction as in the case of PSE_1 and PSE_2 families, where a loop region was predicted to be a transmembrane helix (Refer PSE_1 and PSE_2 topology annotation in CPAfold database). Based on the classification of broken/reentrant type, missing helices were added, and the orientations were corrected. The final topology was then inferred for all families (Figure 2c).

##### iv) Identification of core, scaffold subdomains and inverted repeat units from known structures

Annotations of scaffold and core subdomains were taken from the literature of the Pfam families with known structure. Subdomain annotations were then transferred from the family with an available structure to the family with an unknown structure based on the definition of aligned TM helices described in the previous section (Figure 2c).

##### v) Validation of Broken/Reentrant type of transporters by the positive inside rule

It is well known that the inside loops have an enrichment of positive residues compared to the outside loops of a transmembrane protein[31,56,57]. Therefore, KR bias can be used to identify the orientation of the protein. We counted the number of K (Lysine) and R (Arginine) starting from 10 residues inside the TM helix and up to 25 residues after the helix as this has been shown to contribute to the positive inside rule [58]. The KR-bias is then calculated using the family MSA and comparing the number of KR in the inside and outside loops. Two models were made one representing the broken, and one the reentrant topology. This was then used to confirm the topology of all families/subfamilies. KR bias is calculated for all the helices in the full-length protein, and the expected correct topology would show a higher KR-bias (Figure 2d).

#### 2) Hierarchical clustering

HHsearch program was used to identify the evolutionary relationship of the full-length sequences or repeat units versus the Pfam-A_v32 database. The hit is considered to be N or C repeat based on the alignment with query repeat. If bi-directional pairs of query-hit are obtained, the pair containing the lowest E-value is obtained. Pairs not found by HHsearch were given an E-value higher than the highest E-value observed.

Hierarchical clustering of the log_10_(E-values) was carried out using Hclust in the Heatmap2 program in R using the average cluster method and correlations as the distance. Multiscale Bootstrapping of the hierarchical cluster with 10,000 bootstraps was carried out using the Pvclust program [59] using function hclust in R. AU (Approximately Unbiased) *p*-values for all clusters of original data was computed by multiscale bootstrap resampling.

A phylogenetic tree was generated using IQtree [60] [61] using a multiple sequence alignment of full-length sequences from Clustal Omega [50] as the input. Here, 10 sequences including the representative sequence were randomly considered from topology seed MSA for each family for the construction of MSA. The program also identified the best substitution model (LG+F+G4) that fits the data using the Model Finder program [62]. Bootstrap analysis with 100 replicas were carried out. All the trees were analysed and generated using iTOL[63].

#### 3) Generation of topology alignments between families

MSA-MSA alignment between families obtained in the previous step is converted into pairwise topology alignments. The topology and sequence alignment figure was generated.Some MSA-MSA alignments did not give rise to correctly aligned TM helices, due to uncertainty in introducing long gaps. These cases are accompanied by correctly aligned structure alignments.

#### 4) Sequence motifs in the broken/reentrant helix

The middle helix (Broken/reentrant) of the N and C core subdomain were extracted from the family MSA. Sequence motifs were generated using Weblogo program [64] to access the enriched amino acids in the broken and reentrant helices. Additionally, multiple sequence alignment consisting of the seven helical broken and six helical reentrant fold type was used to study the enrichment of amino acids and assessment of the type of mutations in both the groups.

#### 5) Hydrophobicity (ΔG) and KR bias for broken and reentrant helices

The biological hydrophobicity scale (ΔG) and KR bias were calculated for all the proteins of families. Hydrophobicity of the broken/reentrant helix was calculated using DGpred [65]. KR bias is calculated for the last helix of the N and C core subdomains.

#### 6) Sequence similarity between repeat units

E-values between the N- and C-terminal repeats within the same family are obtained from the full-length alignments using HHsearch program.

#### 7) Structural superposition

Structure superpositions of the pairs of transporters were carried out separately for the core domain and scaffold domain. Structure superpositions were carried out using TMalign [66] and visualised using PyMol [66,67].

## Data availability

All representative sequences, multiple sequence alignments, topology annotations, Core and scaffold domain annotation, KR-bias plots, sequence motifs, full length and repeat level sequence and topology alignments, Structure based classification into fold-types etc. are available in the CPAfold database at cpafold.bioinfo.se for all the families of the CPA/AT superfamily. We also generated homology models of the representative sequence based on the aligned region and the best structural template using the HHpred server [68]. All scripts are available from the github repository https://github.com/ElofssonLab/TMplot

## Acknowledgements

We thank David Drew for valuable input to this manuscript.

## Funding and additional information

We thank the Swedish National Infrastructure for Computing for providing computational resources. This work was supported by a grant VR-NT-2016-03798 from the Swedish National Research Council and a grant from the Kunt and Alice Wallenberg foundation.

